# Identification of Sensory Fiber Types in Mouse Temporomandibular Joint Tissues

**DOI:** 10.1101/2025.05.15.654333

**Authors:** Jessie J. Alfaro, RE-JOIN Consortium Investigators, Armen N. Akopian

## Abstract

Temporomandibular joint (TMJ) disorders (TMJDs) are linked to heightened nerve sensitivity in TMJ tissues. To set the groundwork for investigating the mechanisms governing this increased responsiveness, this study aimed to identify the types of nerves in the retrodiscal tissue (retrodisc), anterior disc, and joint capsule of mouse TMJ using immunohistochemistry (IHC) and reporter mice. The pan-sensory neuronal marker pgp9.5 revealed no nerves in the articular disc but identified approximately 70% unmyelinated and 30% myelinated fibers in other TMJ tissues. Nearly all sensory fibers in the joint capsule and anterior disc were CGRP^+^ peptidergic fibers, while the retrodisc contained about 80% peptidergic fibers. Notably, CGRP^-^/NFH^+^ myelinated non-peptidergic nerves were absent, indicating the absence of non-nociceptive fibers (A-LTMRs) in TMJ tissues. Almost all sensory fibers in the joint capsule and anterior disc were Htr3a^+^, with the retrodisc containing 60-70% Htr3a^+^ fibers. Additionally, TMJ tissues had minimal to no (<5%) MrgprD^+^, MrgprA3^+^, MrgprC11^+^, somatostatin^+^, or parvalbumin^+^ fibers, except for the retrodisc, which had about 20% Mrgpr^+^ fibers. Excluding articular discs, TMJ tissues were highly vascularized, with blood vessels surrounded by both sensory and sympathetic (TH+) nerves. Overall, TMJ tissues were predominantly innervated by peptidergic fibers, with a minor presence of other non-peptidergic nociceptors.

## Introduction

Temporomandibular joint (TMJ) disorders (TMJDs) comprise a group of conditions that impair TMJ function, with both painful and non-painful forms ^1,2^. Painful TMJDs affect around 40% of patients—approximately 30 million people in the U.S.—and 15% of these individuals experience chronic pain ^2–4^. This pain is often the most challenging symptom to manage^2^. Conservative treatments (e.g., soft diet, physical therapy, NSAIDs) frequently provide insufficient relief, prompting the use of more invasive options like arthrocentesis or total joint replacement ^1,5^. Treatment selection is typically based on clinical criteria such as the Wilkes classification ^1,5^.

A major barrier to effective pain management in TMJD is the limited understanding of its underlying molecular mechanisms. This knowledge gap underscores the need to characterize TMJ sensory innervation. The TMJ is innervated by the auriculotemporal (ACN), masseteric (MN), and anterior deep TMJ nerves—branches of the mandibular division (V3) of the trigeminal nerve^6^.

Prior studies indicate a predominance of C– and A-fiber nociceptors in TMJ tissues, with TMJ afferents expressing neuropeptides such as calcitonin gene-related peptide (CGRP), substance P (SP), neuropeptide Y (NPY), and nitric oxide synthase (NOS)^7–10^. CGRP-positive nerves were found in the capsule and the synovial membrane, but not in the articular disc of sheep^11,12^. *In vivo* extracellular recordings from TMJ structures – ATN single nerve preparations demonstrated that TMJ nerves were classified into the following four subtypes: Adelta-high-threshold mechanonociceptor (Aδ-HTMR) (12.1%), Adelta-polymodal nociceptor (Aδ-POLY) (36.4%), C-HTMR (12.1%), and C-POLY (39.4%)^13,14^. Overall, despite accumulated data on sensory neurons innervating TMJ, there is limited information regarding the precise make-up of neuronal subtypes innervating the TMJ tissues, such as the joint capsule and associated synovium, articular disc, anterior disc, and retrodisc.

The development of single-cell RNA sequencing technology opened avenues for precise identifications of sensory nerve subtypes in tissues using IHC with well-defined antibody markers and reporter mouse lines highlighting expressions of sensory neuronal markers^20^. Accordingly, the main aim of this study was to identify nerve subtypes in each TMJ tissue using IHC and reporter mouse lines.

## Results

### TMJ tissues examined in this study

TMJ tissues exhibit distinct functions and pathophysiological profiles, and are believed to undergo unique gene expression changes and cellular plasticity during TMJD. Therefore, this study focused on characterizing the innervation and vascularization of individual TMJ structures ^9,15,16^. We generated serial cryosections that sequentially passed through the mandible, temporal bone, and a thin layer of the joint capsule, which contains synovial tissue in its deeper layers (*Fig 1A*). As the sections progressed, they encompassed the lateral pterygoid muscle (LPM), condyle, anterior disc, articular disc, and retrodiscal tissue (aka retrodisc) (*Fig 1B*). Innervation and vascularization were analyzed in the joint capsule (*Fig. 1C*), as well as in the anterior disc and retrodisc (*Fig. 1D*). These features were not assessed in the articular disc due to inconclusive evidence of nerves or blood vessels in this region. It remains unclear whether the neurovascular elements observed in the anterior disc and retrodisc extend into and are functionally present within the articular disc itself (*Figs. 1E, 1F*).

**Figure 1.**
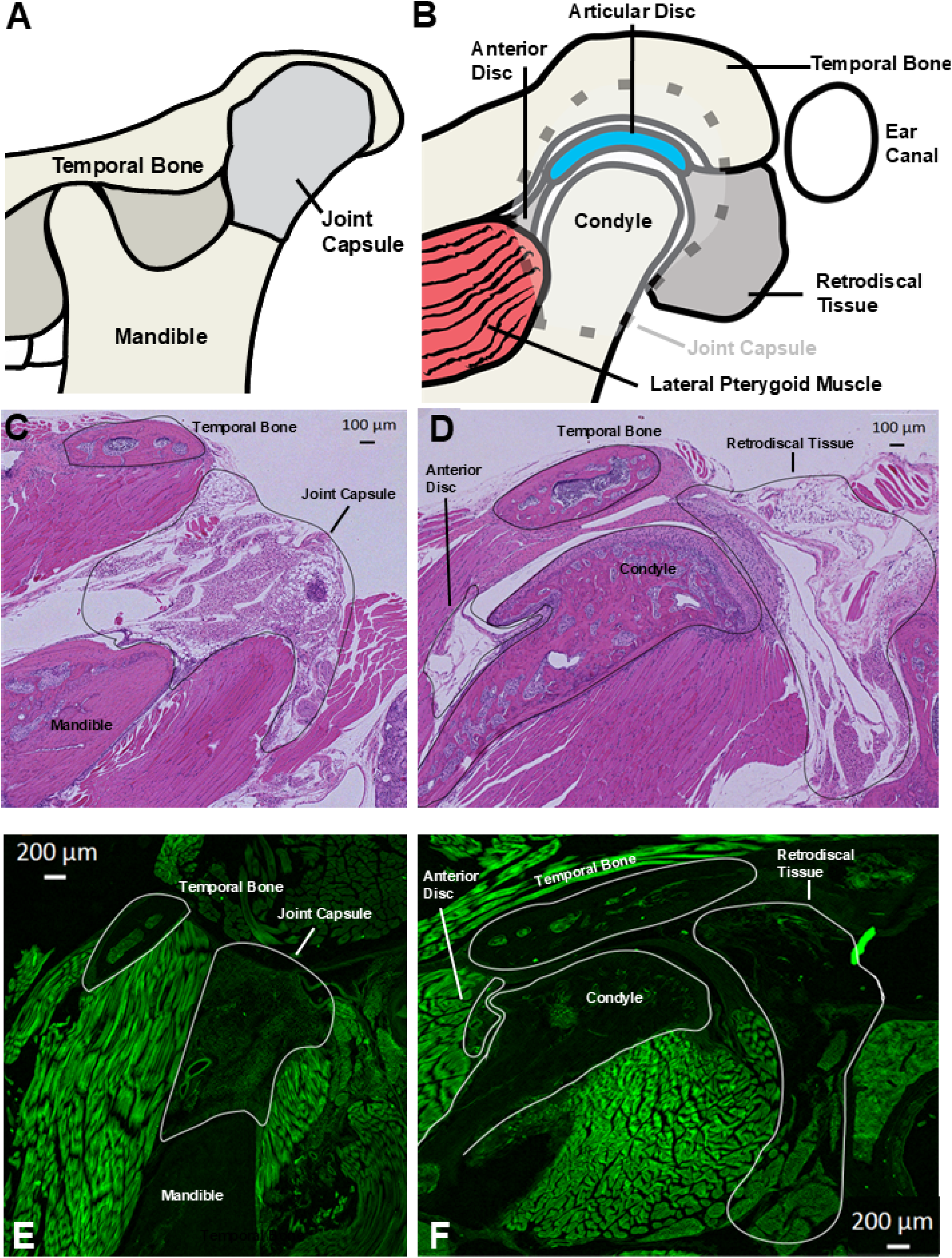
Schematic for TMJ tissues. (**A**) Schematic of TMJ tissues on the initial serial sections. (**B**) Schematic of TMJ tissues in dipper serial sections. Dashed line shows position of the joint capsule overlaying TMJ deeper tissues, such as the anterior disc, articulate disc and retrodiscal tissue. (**C**) The joint capsula as well as temporal bone and mandible are outlined in an H&E paraffin section. (**D**) The anterior disc and retrodiscal tissue as well as condyle and temporal bone are outlined in an H&E paraffin section. (**E**) The joint capsula is outlined in a cryo-section. (**F**) The anterior disc and retrodiscal tissue are outlined in a cryo-section. TMJ tissues are labeled and outlined on the panels A-F.

### Myelinated and unmyelinated sensory afferents in TMJ tissues

All sensory nerves were identified using the pan-neuronal marker pgp9.5, while myelinated fibers were specifically labeled with NFH antibodies ^17,18^. In the joint capsule, anterior disc, and retrodisc, approximately 30% of sensory afferents were myelinated (pgp9.5⁺/NFH⁺), while the majority—around 70%—were unmyelinated (pgp9.5⁺/NFH⁻) (*Figs. 2A, 2B*). This pattern indicates significantly higher innervation by unmyelinated fibers across these TMJ tissues. Ovell, in contrast to masticatory muscles, which typically exhibit a higher proportion of myelinated fibers^17,18^, TMJ structures were predominantly innervated by unmyelinated sensory nerves. Despite this predominance, the relative proportions of myelinated and unmyelinated fibers were consistent across the joint capsule, anterior disc, and retrodisc.

**Figure 2.**
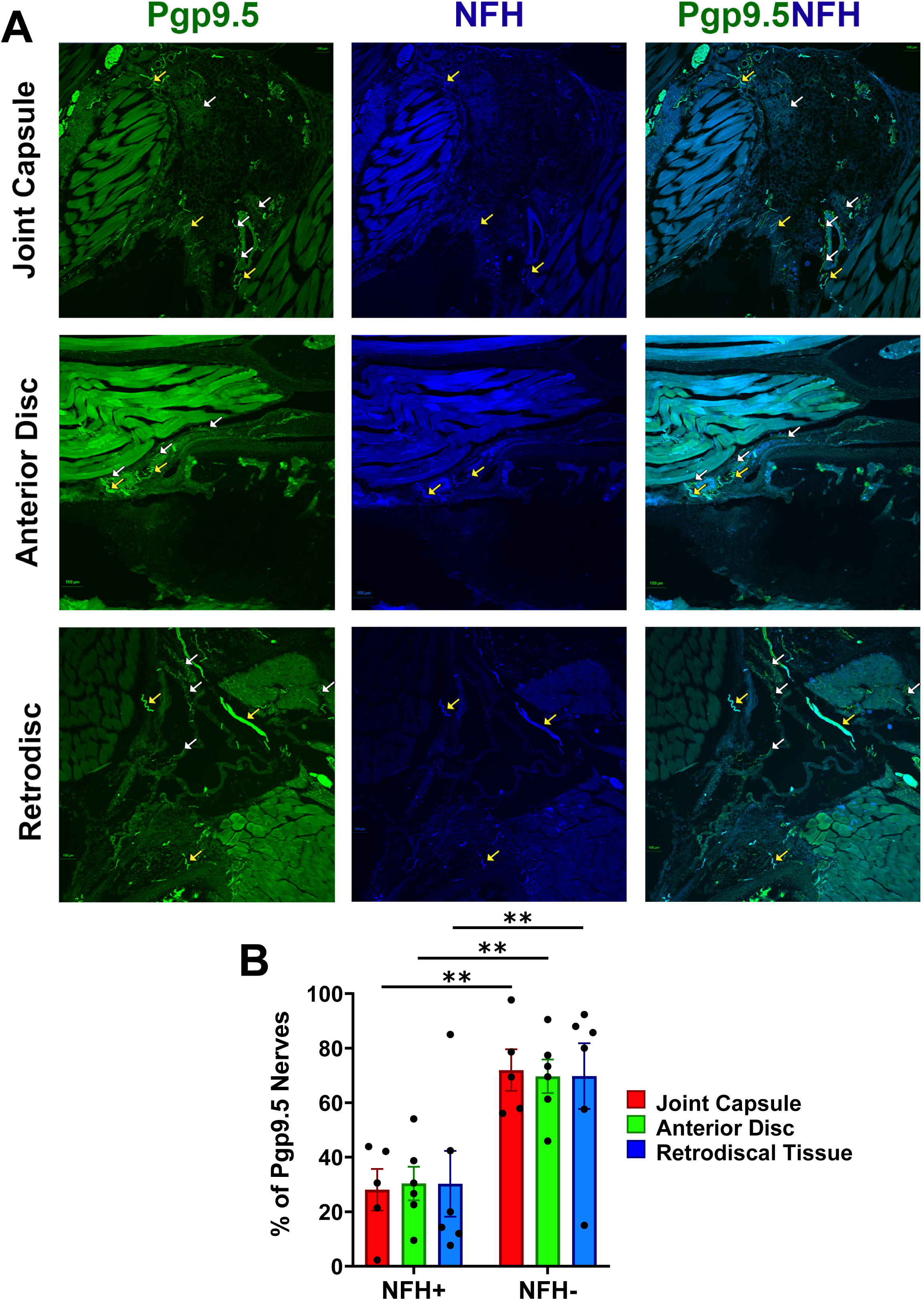
Pgp9.5 and NFH-positive fibers in the joint capsule, anterior disc and retrodiscal tissues. (**A**) The *left column* shows pgp9.5-positive fibers (all sensory fibers), the *middle column* shows NFH-positive (A-fibers) fibers and the *right column* shows co-expressions of pgp9.5 and NFH-positive fibers in the mouse joint capsule, anterior disc and retrodiscal tissues. Pictures from the joint capsule, anterior disc and retrodiscal tissues as well as antibodies used and corresponding colors are indicated. Yellow arrows mark pgp9.5^+^/NFH^+^ fibers, and white arrows show pgp9.5^+^/NFH^-^ nerves. (**B**) Percentages of pgp9.5^+^/NFH^+^ and pgp9.5^+^/NFH^-^ nerves in the joint capsule, anterior disc and retrodiscal tissues. Statistics is 2-way ANOVA Bonferroni’s pot-hoc test (** p<0.01; n=5-6).

### Peptidergic and Htr3a-positive sensory afferents in TMJ tissues

We next assessed the presence of CGRP⁺ peptidergic nerves in TMJ tissues. The joint capsule and anterior disc were almost entirely populated by peptidergic sensory fibers (*Figs. 3A, 3B*), while the retrodiscal tissue contained approximately 80% CGRP⁺/pgp9.5⁺ nerves (*Figs. 3A, 3B*). Further analysis using CGRP reporter mice and co-labeling with CGRP and NFH antibodies confirmed that roughly 70% of peptidergic fibers were myelinated, with the remaining 30% unmyelinated (*Figs. 4A, 4B*). Notably, TMJ tissues showed minimal to no (<5%) CGRP⁻/NFH⁺ fibers—non-peptidergic myelinated sensory nerves typically classified as non-nociceptive Aβ low-threshold mechanoreceptors (A-LTMRs) ^12,19,20^.

**Figure 3.**
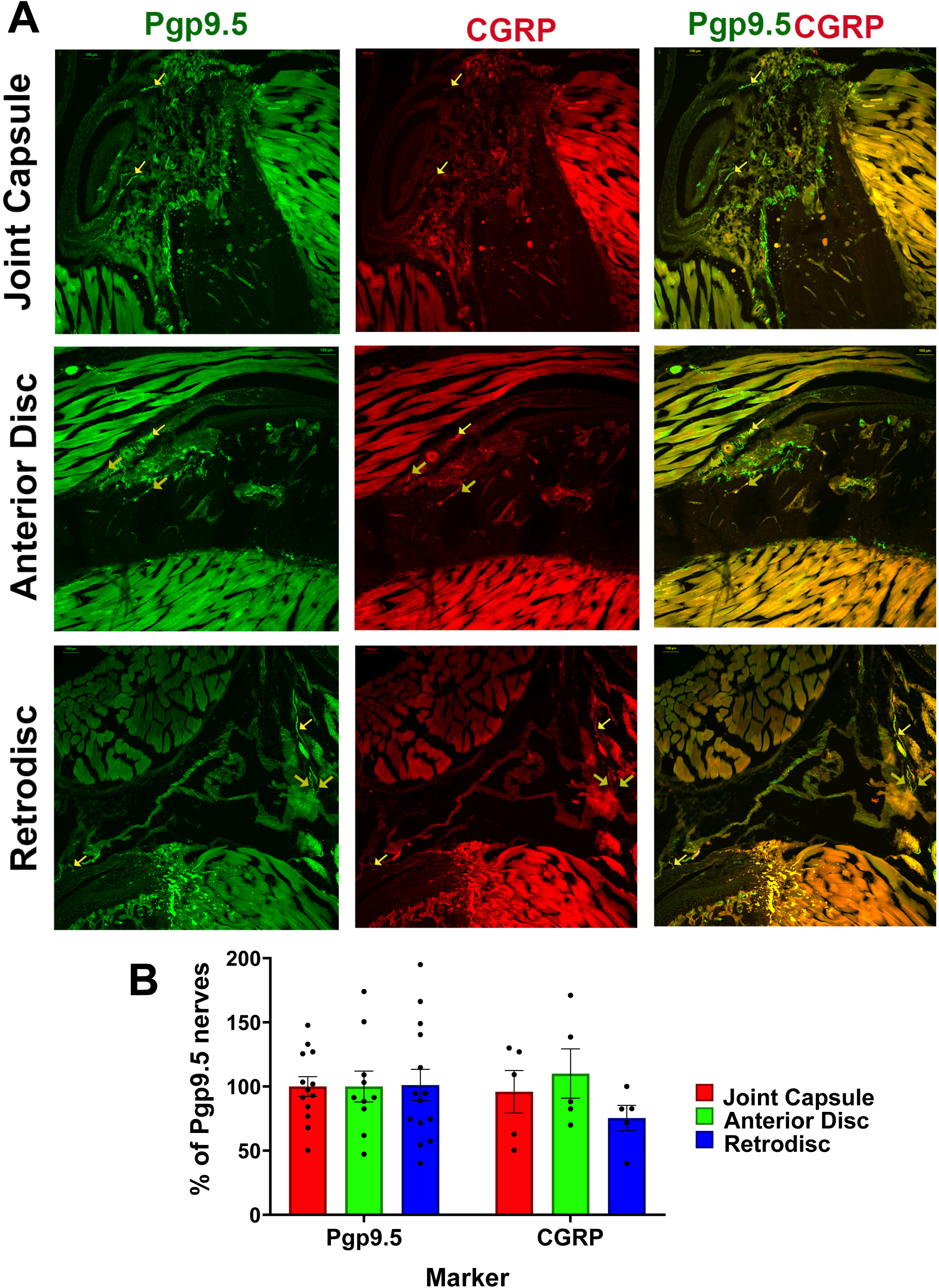
Peptidergic fibers in the joint capsule, anterior disc and retrodiscal tissues. (**A**) The *left column* shows pgp9.5-positive fibers (all sensory fibers), the *middle column* shows CGRP-positive (peptidergic fibers) fibers in CGRP/tdTom mouse and the *right column* shows co-expressions of CGRP and pgp9.5-positive fibers in the mouse joint capsule, anterior disc and retrodiscal tissues. Pictures from the joint capsule, anterior disc and retrodiscal tissues as well as antibodies used and corresponding colors are indicated. Yellow arrows mark pgp9.5^+^/CGRP^+^ fibers. (**B**) Percentages of pgp9.5^+^ and CGRP^+^ nerves relative to pgp9.5^+^ in a selected section for the joint capsule, anterior disc and retrodiscal tissues. Statistics is 2-way ANOVA Bonferroni’s pot-hoc test (n=5-13).

**Figure 4.**
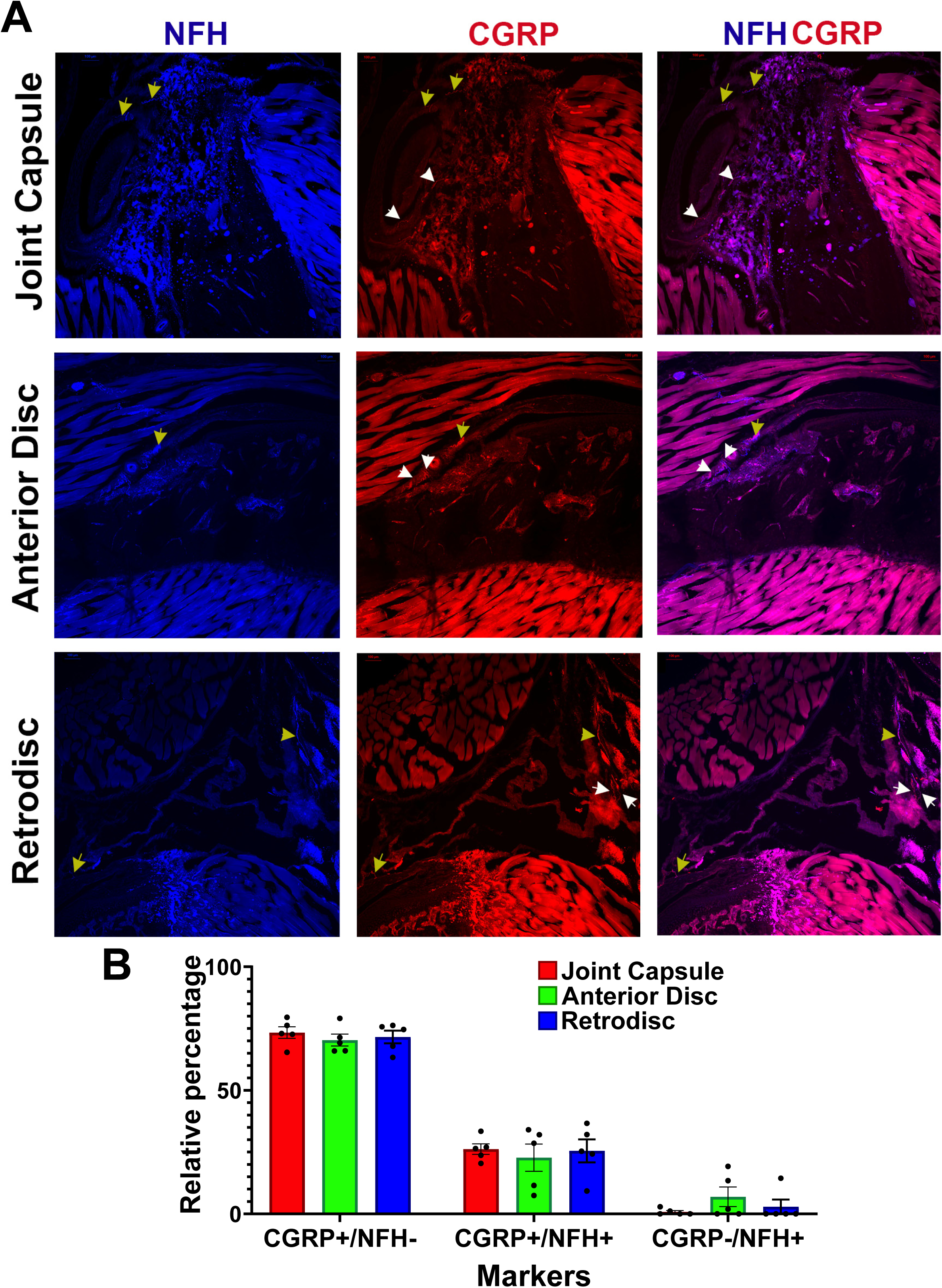
Peptidergic myelinated and non-myelinated fibers in the joint capsule, anterior disc and retrodiscal tissues. (**A**) The *left column* shows NFH-positive fibers (A-fibers), the *middle column* shows CGRP-positive (peptidergic fibers) fibers in CGRP/tdTom mouse and the *right column* shows co-expressions of CGRP and NFH-positive fibers in the mouse joint capsule, anterior disc and retrodiscal tissues. Pictures from the joint capsule, anterior disc and retrodiscal tissues as well as antibodies used and corresponding colors are indicated. Yellow arrows mark CGRP^+^/NFH^+^ fibers and white arrows show CGRP^+^/NFH^-^ nerves. (**B**) Relative percentages for CGRP^+^/NFH^-^, CGRP^+^/NFH^-^ and CGRP^-^/NFH^+^ nerves in the joint capsule, anterior disc and retrodiscal tissues. Statistics is 2-way ANOVA Bonferroni’s pot-hoc test (n=5).

Given that several peptidergic fiber subtypes express Htr3a ^18^, we examined this further using Htr3a/tdTomato reporter mice. In the joint capsule and anterior disc, Htr3a⁺ fibers almost completely overlapped with pgp9.5⁺ nerves, whereas the retrodisc showed only about 50% overlap (*Figs. 5A, 5B*). In the joint capsule and anterior disc, Htr3a⁺ fibers almost completely overlapped with pgp9.5⁺ nerves, whereas the retrodisc showed only about 50% overlap (*Figs. 5A, 5B*). As expected, approximately 70% of Htr3a⁺ fibers were unmyelinated, with the remainder being myelinated; no Htr3a⁻/NFH⁺ fibers were detected (Fig. 5C) ^18^. In summary, nearly all sensory nerves in the joint capsule and anterior disc are peptidergic and co-express Htr3a. In contrast, the retrodisc contains a distinct subset (∼20%) of non-peptidergic, Htr3a⁻ fibers^18^, along with some peptidergic fibers that are likely Htr3a⁻.

**Figure 5.**
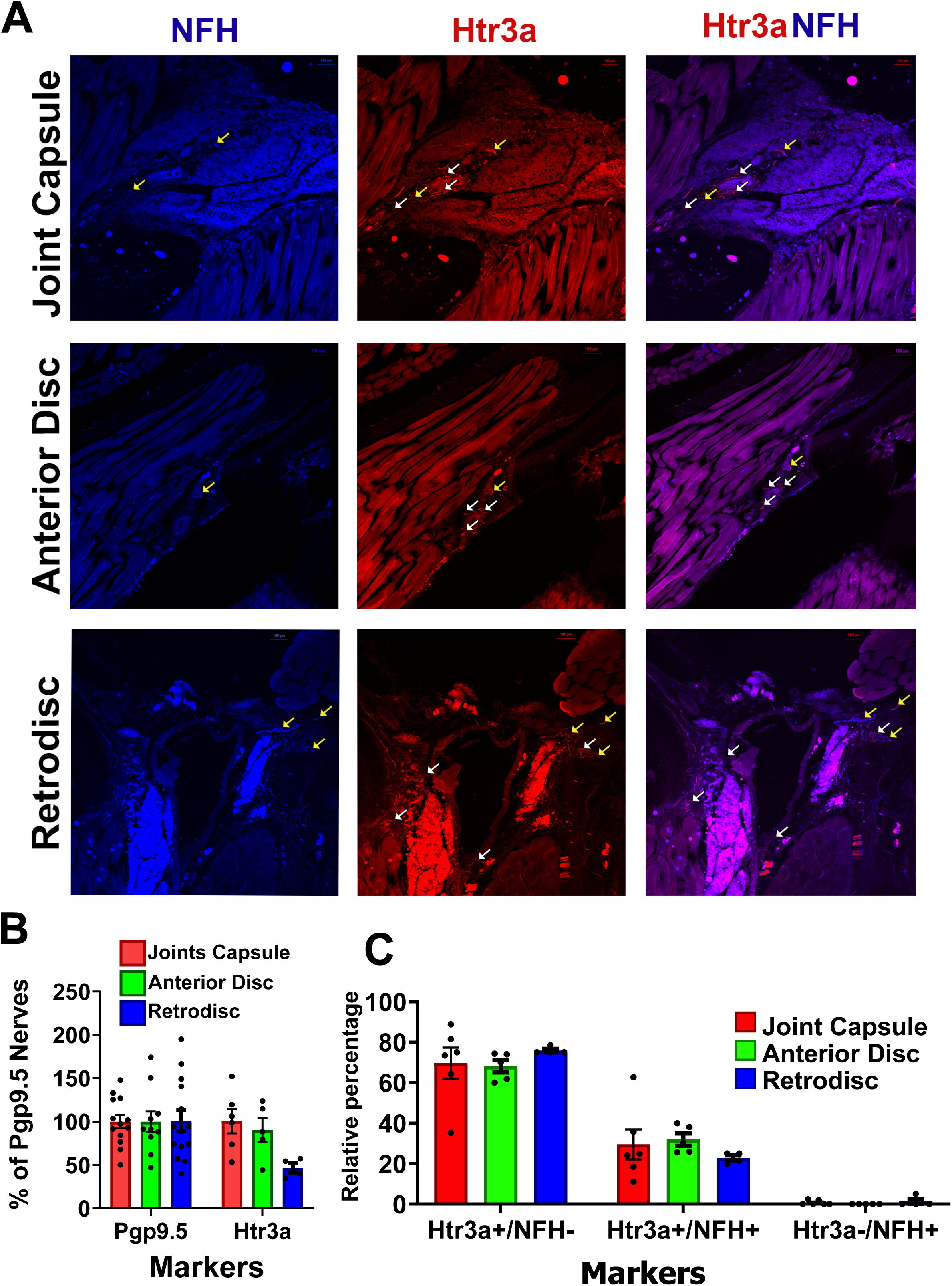
Htr3a-positive fibers in the joint capsule, anterior disc and retrodiscal tissues. (**A**) The *left column* shows NFH-positive fibers (A-fibers), the *middle column* shows Htr3a-positive fibers in Htr3a/tdTom mouse and the *right column* shows co-expressions of Htr3a and NFH-positive fibers in the mouse joint capsule, anterior disc and retrodiscal tissues. Pictures from the joint capsule, anterior disc and retrodiscal tissues as well as antibodies used and corresponding colors are indicated. Yellow arrows mark Htr3a^+^/NFH^+^ fibers, and white arrows show Htr3a^+^/NFH^-^ nerves. (**B**) Percentages of pgp9.5^+^ and Htr3a^+^ nerves relatively to pgp9.5^+^ in a selected section for the joint capsule, anterior disc and retrodiscal tissues. Statistics is 2-way ANOVA Bonferroni’s pot-hoc test (n=5-13). (**C**) Relative percentages for Htr3a^+^/NFH^-^, Htr3a^+^/NFH^-^ and Htr3a^-^/NFH^+^ nerves in the joint capsule, anterior disc and retrodiscal tissues. Statistics is 2-way ANOVA Bonferroni’s pot-hoc test (n=5-6).

### Presence of non-peptidergic – MrgprD, MrgprA3, MrgprC11, somatostatin and parvalbumin *-sensory afferents in TMJ tissues*

These findings suggest that TMJ tissues are predominantly innervated by peptidergic sensory fibers, with widespread expression of Htr3a. To further validate this, we examined the presence of non-peptidergic nerve subtypes, focusing on MrgprD⁺ fibers, which represent the major class of non-peptidergic neurons (NP-1 group) in the dorsal root ganglia (DRG) and trigeminal ganglia (TG) ^21^. Using MrgprD/tdTomato reporter mice, we found that MrgprD⁺ fibers were scarce in the joint capsule and anterior disc, comprising less than 5% of the total sensory innervation. In contrast, the retrodisc showed a modestly higher presence, with approximately 12% MrgprD⁺ fibers (*Figs. 6A, 6B*). As a positive control, facial skin displayed dense MrgprD⁺ innervation, confirming the sensitivity of our detection method (*Suppl. Fig. 1*). These results reinforce the conclusion that TMJ tissues are mainly innervated by peptidergic neurons, while non-peptidergic MrgprD⁺ fibers are limited, particularly outside the retrodisc.

**Figure 6.**
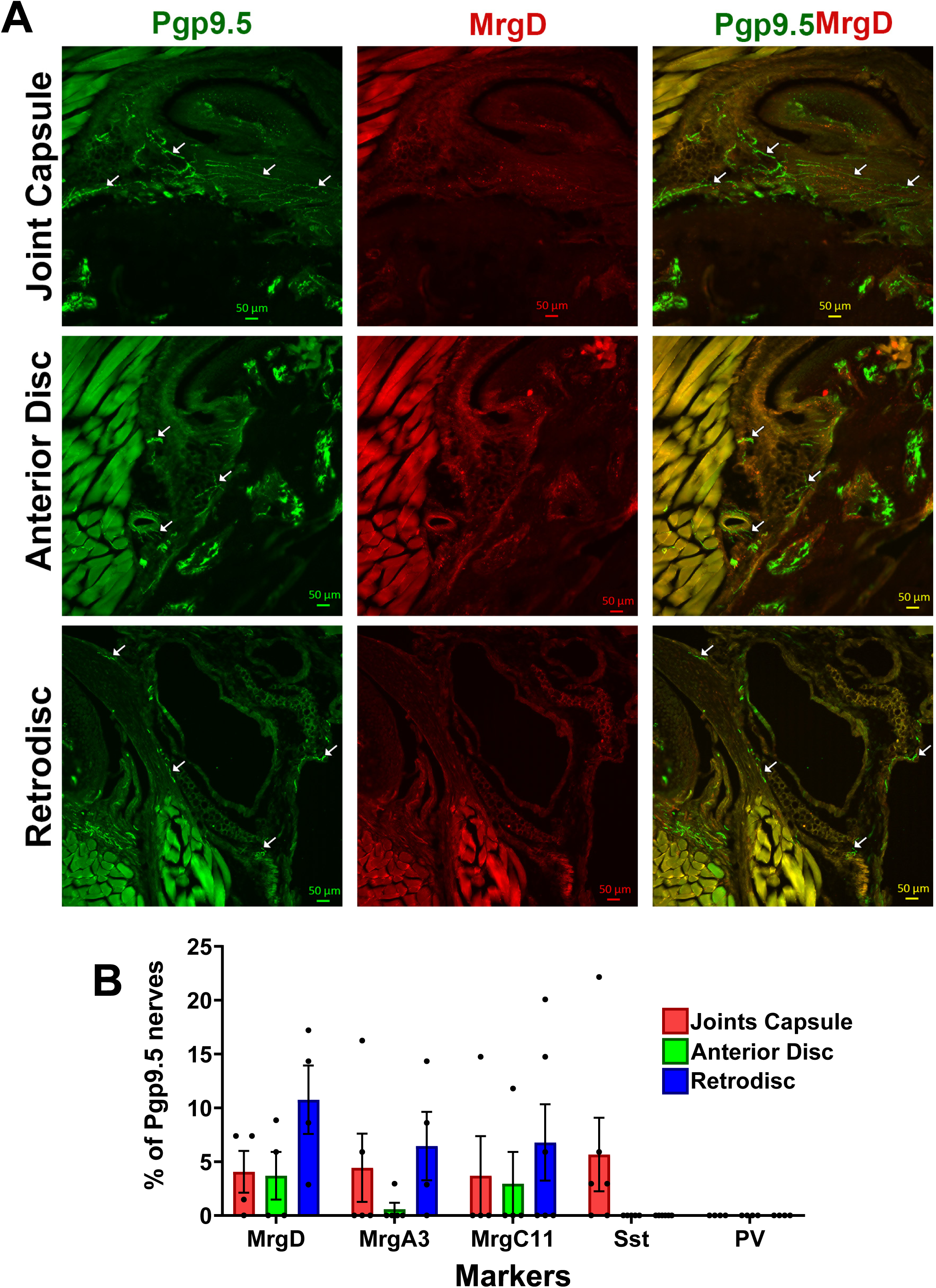
MrgprD-positive fibers in the joint capsule, anterior disc and retrodiscal tissues. (**A**) The *left column* shows pgp9.5-positive fibers (all sensory fibers), the *middle column* shows MrgprD-positive fibers in MrgprD/tdTom mouse and the *right column* shows co-expressions of pgp9.5 and MrgprD-positive fibers in the mouse joint capsule, anterior disc and retrodiscal tissues. Pictures from the joint capsule, anterior disc and retrodiscal tissues as well as antibodies used and corresponding colors are indicated. White arrows show pgp9.5^+^/MrgprD^-^ nerves. (**B**) Percentages of MrgprD^+^, MrgprA3^+^, MrgprC11^+^, Sst^+^ and PV^+^ nerves relatively to numbers of pgp9.5^+^ nerves for the joint capsule, anterior disc and retrodiscal tissues. Statistics is 2-way ANOVA Bonferroni’s pot-hoc test (n=4-5).

Another subset of non-peptidergic fibers, known as the NP-2 group, is marked by MrgprA3⁺ expression ^19,21^. However, some studies have shown that MrgprA3⁺ neurons are heterogeneous, comprising both peptidergic and non-peptidergic subtypes ^22,23^. In MrgprA3/tdTomato reporter mice, MrgprA3⁺ nerves were detected only sporadically in the joint capsule and retrodiscal tissues, with none observed in the anterior disc (*Figs. 6B, 7*). A broader subset, marked by MrgprC11⁺, includes certain TrpV1⁺ peptidergic neurons ^24^. In MrgprC11/tdTomato reporter mice, TMJ tissues displayed only a small number of MrgprC11⁺ nerves (*Figs. 6B, 8*). As with MrgprD⁺ fibers, facial skin—used as a positive control—showed significantly greater innervation by both MrgprA3⁺ and MrgprC11⁺ fibers (*Suppl. Fig. 1*). These results further support the conclusion that TMJ tissues contain very few non-peptidergic sensory fibers, with only limited contributions from MrgprA3⁺ and MrgprC11⁺ subtypes.

**Figure 7.**
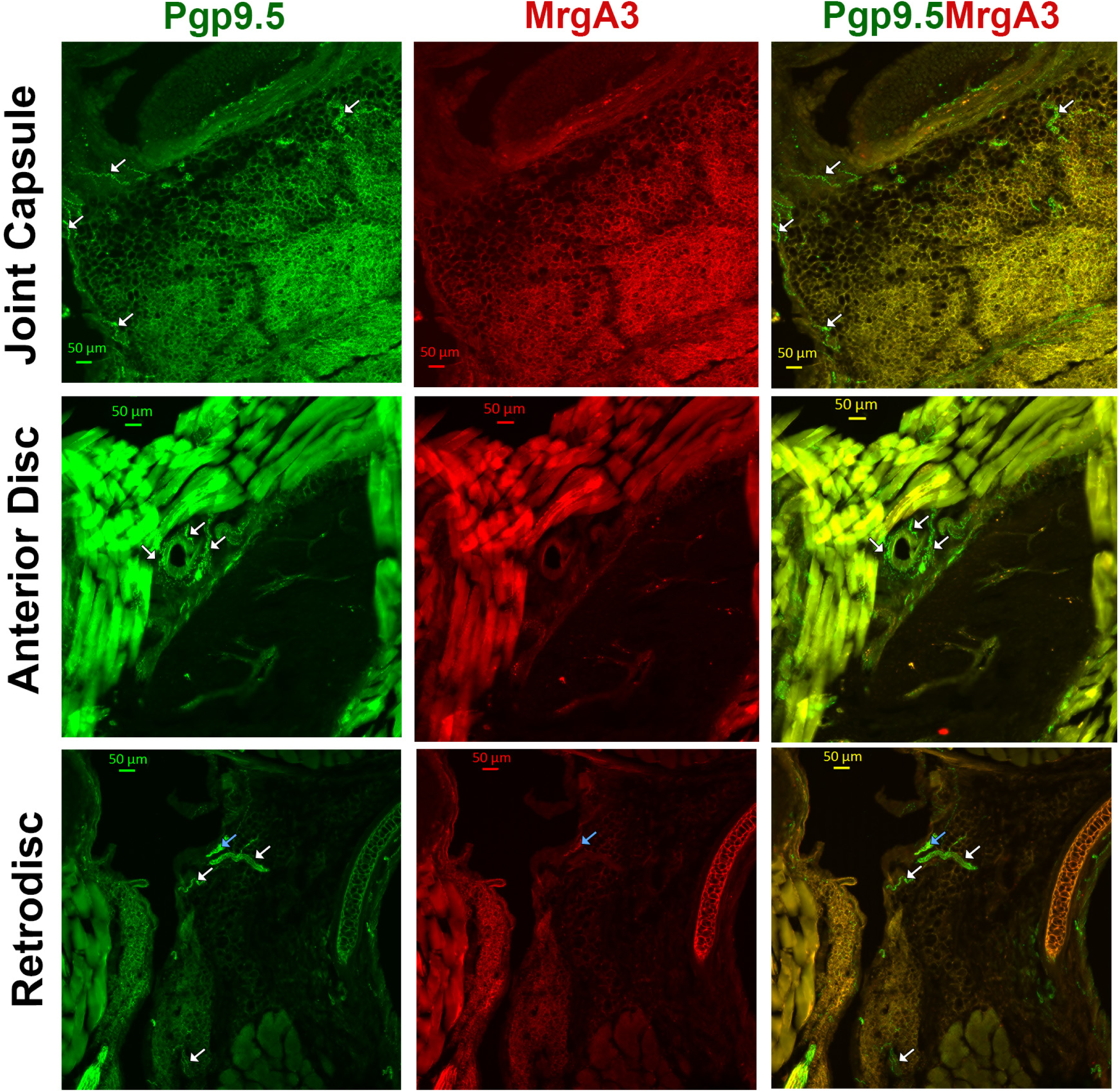
MrgprA3-positive fibers in the joint capsule, anterior disc and retrodiscal tissues. *The left column* shows pgp9.5-positive fibers (all sensory fibers), the *middle column* shows MrgprA3-positive fibers in MrgprA3/tdTom mouse and the *right column* shows co-expressions of pgp9.5 and MrgprA3-positive fibers in the mouse joint capsule, anterior disc and retrodiscal tissues. Pictures from the joint capsule, anterior disc and retrodiscal tissues as well as antibodies used and corresponding colors are indicated. Blue arrow marks a pgp9.5^+^/MrgprA3^+^ fiber, and white arrows show pgp9.5^+^/MrgprA3^-^ nerves.

**Figure 8.**
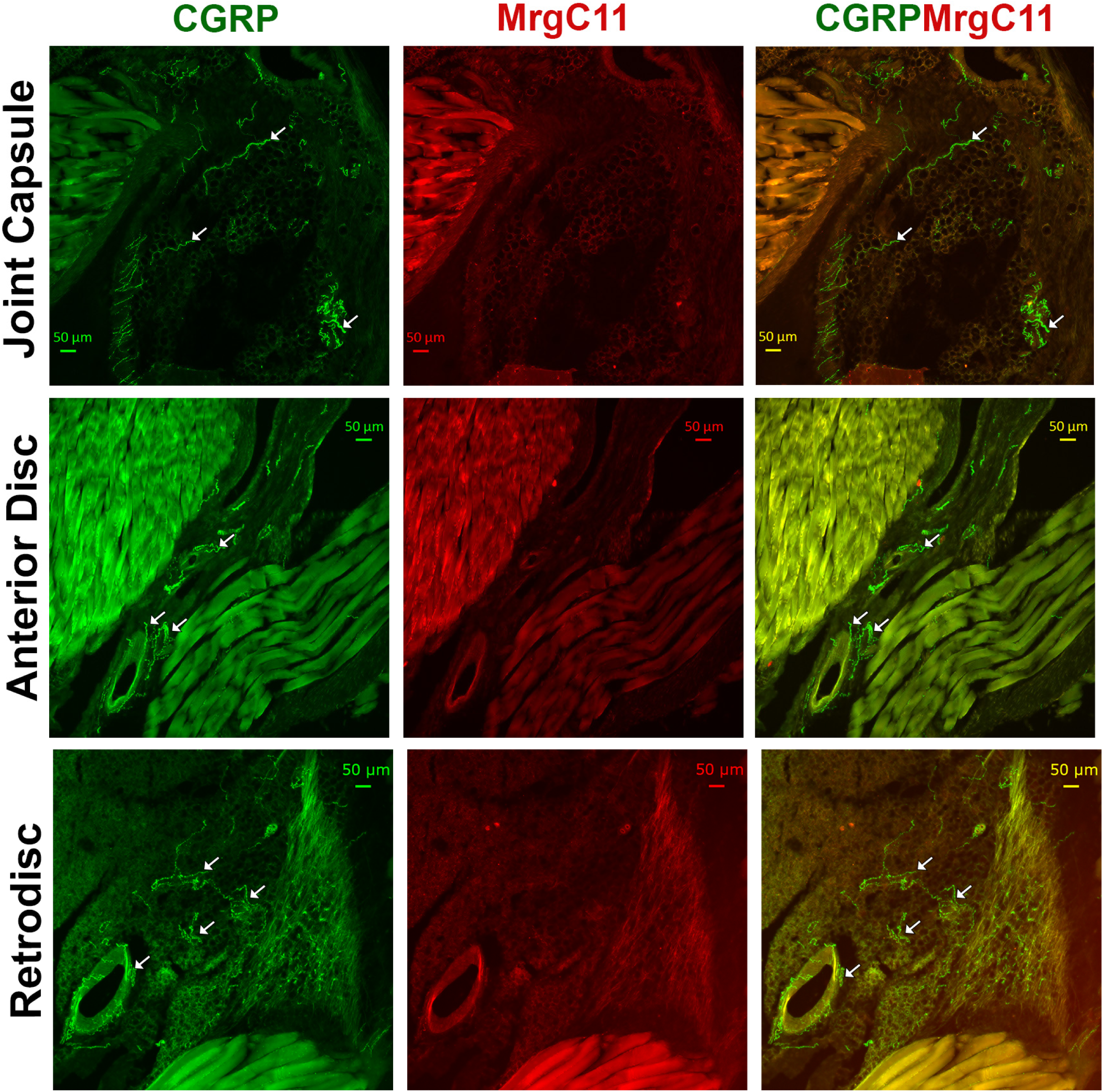
MrgprC11-positive peptidergic fibers in the joint capsule, anterior disc and retrodiscal tissues. The *left column* shows CGRP-positive fibers (peptidergic fibers), the *middle column* shows MrgprC11-positive fibers in MrgprC11/tdTom mouse and the *right column* shows co-expressions of CGRP and MrgprA3-positive fibers in the mouse joint capsule, anterior disc and retrodiscal tissues. Pictures from the joint capsule, anterior disc and retrodiscal tissues as well as antibodies used and corresponding colors are indicated. White arrows show CGRP^+^/MrgprC11^-^ nerves.

The final group of non-peptidergic nociceptors in the DRG and TG is marked by somatostatin (Sst⁺), also referred to as the NP-3 cluster ^12,21^. In Sst/tdTomato reporter mice, Sst⁺ fibers were not detected in the anterior disc or retrodisc, while the joint capsule contained a small proportion— approximately 6%—of Sst⁺ nerves (*Figs. 6A, 9*). The dura mater was used as a positive control and showed robust Sst⁺ innervation (*Suppl. Fig. 1*), validating the specificity of our detection. Overall, TMJ tissues exhibited very low levels of non-peptidergic innervation. Among the TMJ regions analyzed, the retrodisc consistently contained the highest proportion of non-peptidergic sensory fibers.

**Figure 9.**
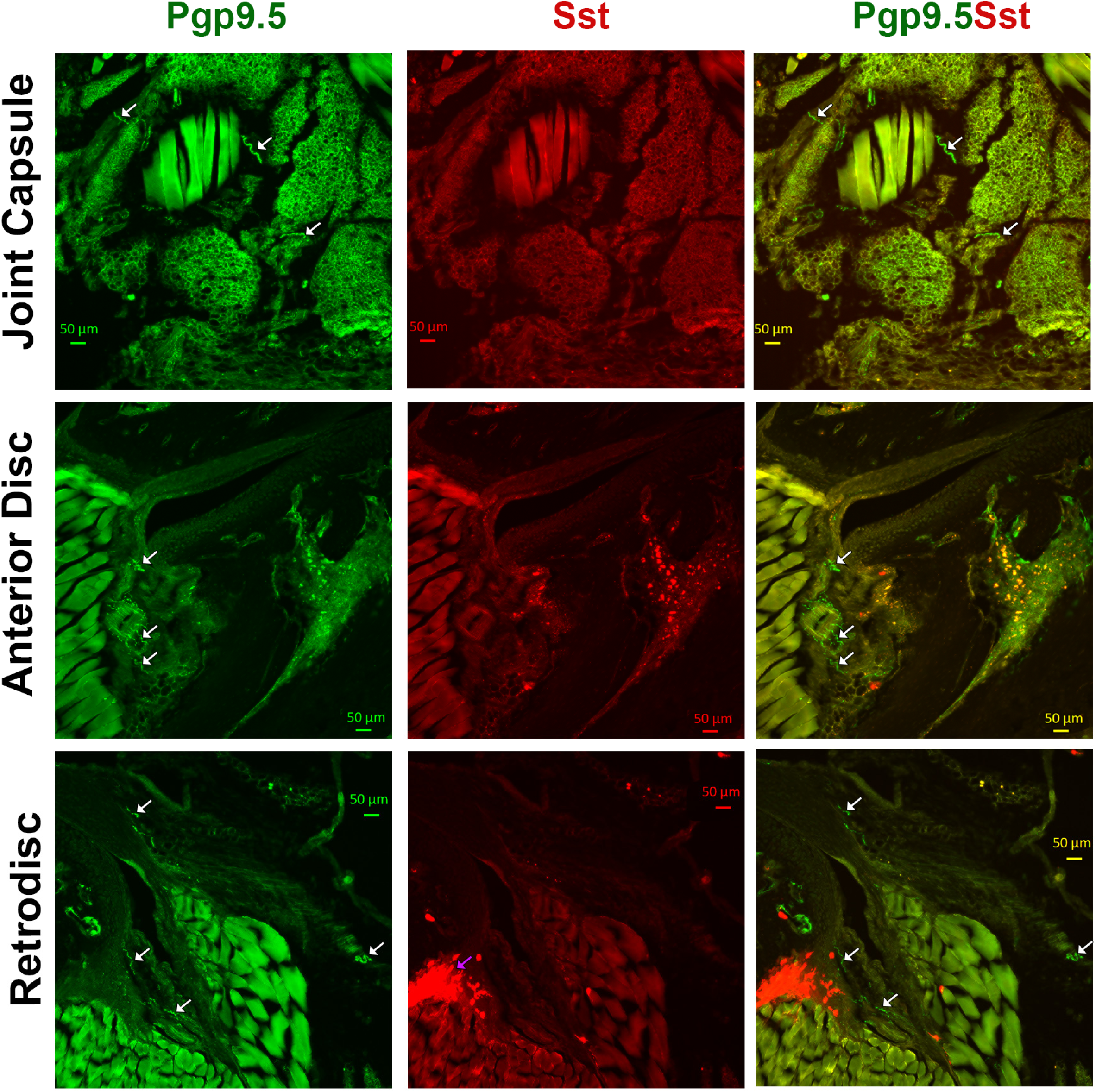
Somatostatin (Sst)-positive fibers in the joint capsule, anterior disc and retrodiscal tissues. The *left column* shows pgp9.5-positive fibers (all sensory fibers), the *middle column* shows Sst-positive fibers in Sst/tdTom mouse and the *right column* shows co-expressions of pgp9.5 and Sst-positive fibers in the mouse joint capsule, anterior disc and retrodiscal tissues. Pictures from the joint capsule, anterior disc and retrodiscal tissues as well as antibodies used and corresponding colors are indicated. White arrows show pgp9.5^+^/Sst^-^ nerves and pink arrow marks a non-neuronal Sst^+^ cells in the retrodiscal tissue.

Non-peptidergic, non-nociceptive nerves are typically classified as low-threshold mechanoreceptors (LTMRs), which include C-, Aδ-, and Aβ-LTMR subtypes ^19,21^. These subtypes are commonly marked by expression of vGlut3 or tyrosine hydroxylase (TH) for C-LTMRs, TrkB for Aδ-LTMRs, and TrkC or parvalbumin (PV) for Aβ-LTMRs ^21^. While both mouse and primate trigeminal ganglia (TGs) contain few TH⁺ neurons, they show a high prevalence of PV⁺ neurons^17,18^. Similarly, facial skin—used here as a positive control—exhibited abundant PV⁺ nerve fibers (*Suppl. Fig. 1*). To assess the presence of LTMRs in TMJ tissues, we analyzed PV/tdTomato reporter mice. No PV⁺ nerve fibers were detected in the joint capsule, anterior disc, or retrodisc (*Figs. 6B, 10*). In summary, TMJ tissues contain very few—if any—non-peptidergic nociceptors or non-nociceptive mechanoreceptors. Among the TMJ regions examined, the retrodisc consistently exhibited the highest proportion of non-peptidergic nociceptors, accounting for approximately 15– 20% of the total innervation.

**Figure 10.**
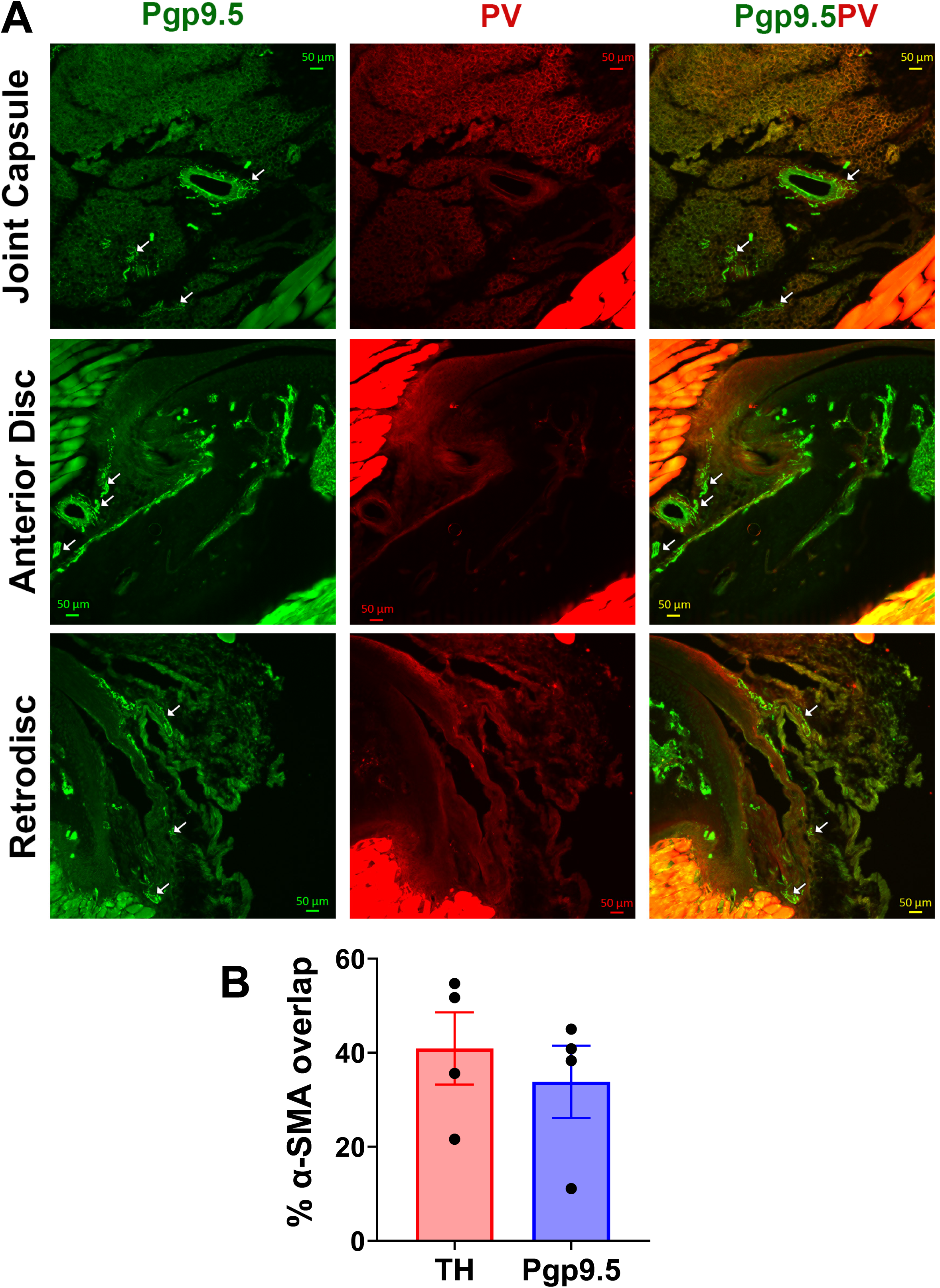
Parvalbumin (PV)-positive fibers in the joint capsule, anterior disc and retrodiscal tissues. (**A**) The *left column* shows pgp9.5-positive fibers (all sensory fibers), the *middle column* shows PV-positive fibers in PV/tdTom mouse and the *right column* shows co-expressions of pgp9.5 and PV-positive fibers in the mouse joint capsule, anterior disc and retrodiscal tissues. Pictures from the joint capsule, anterior disc and retrodiscal tissues as well as antibodies used and corresponding colors are indicated. White arrows show pgp9.5^+^/PV^-^ nerves. (**B**) Percentages of pgp9.5^+^ and tyrosine hydroxylase (TH)-positive nerves in a vicinity of the blood vessels (α-SMA^+^) in the joint capsule, anterior disc and retrodiscal tissues. Statistics is 1-way ANOVA Bonferroni’s pot-hoc test (n=4).

### Innervation of vasculature in TMJ tissues

Using α-SMA as a marker for vascular smooth muscle, we clearly identified blood vessels in all TMJ tissues except the articular disc (*Figs. 10B, 11, 12*). To determine whether these vessels were innervated by sensory or sympathetic nerves, we used pgp9.5 as a pan-sensory nerve marker and tyrosine hydroxylase (TH) to label sympathetic fibers. Both sensory and sympathetic nerves were detected in the joint capsule, anterior disc, and retrodisc, but not in the articular disc (*Figs. 11, 12*). Nerves were considered to innervate blood vessels if they overlapped or were located within 50 µm of α-SMA⁺ vessels. Approximately 35% of all pgp9.5⁺ sensory fibers were found in close association with blood vessels (*Figs. 10B, 11*), while sympathetic TH⁺ fibers showed a slightly higher association at around 40% (*Figs. 10B, 12*). Overall, these findings indicate that TMJ tissues—excluding the articular disc—are well vascularized and receive comparable innervation from both sensory and sympathetic nerves. The joint capsule, anterior disc, and retrodisc displayed similar patterns of neurovascular pattern.

**Figure 11.**
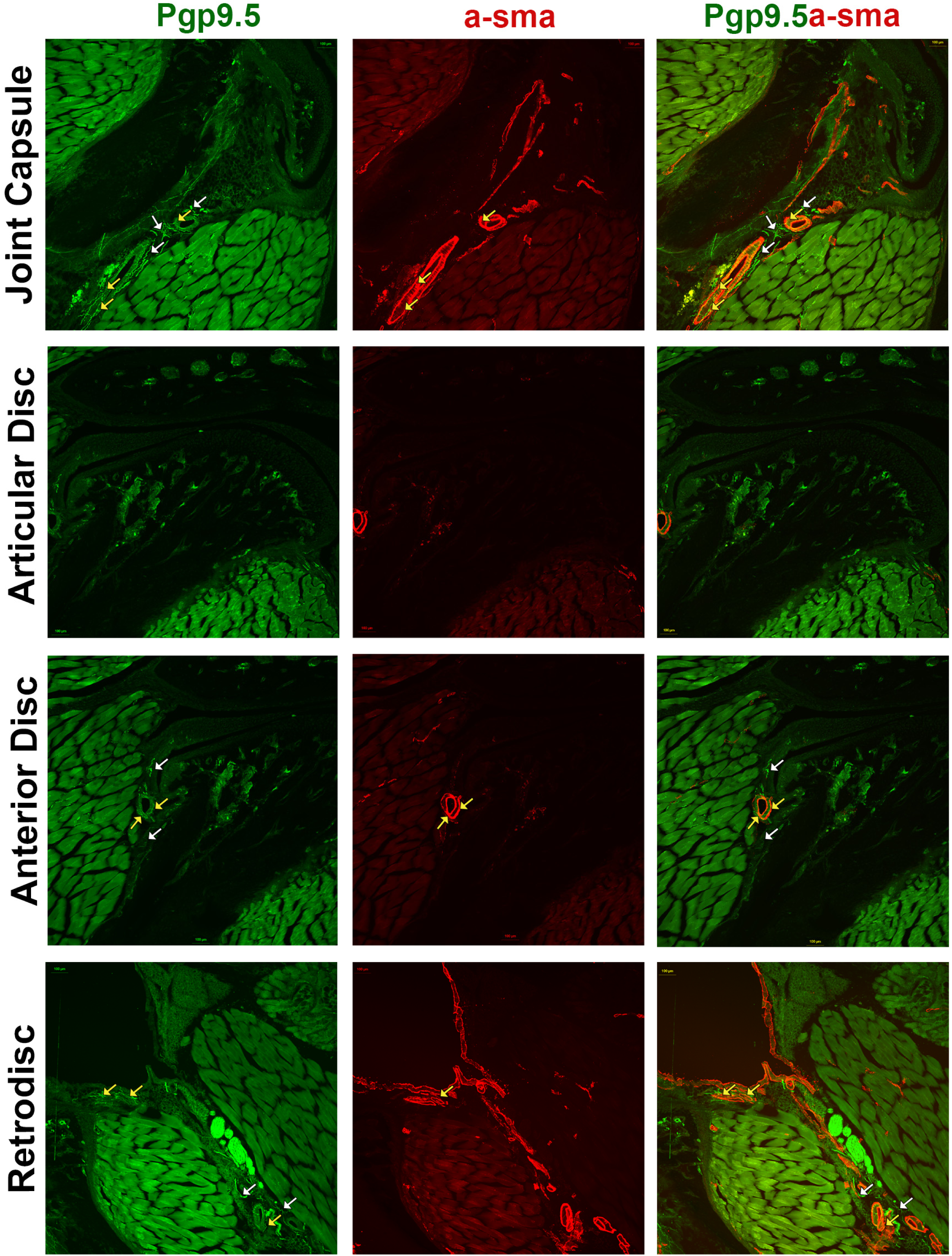
Innervation of blood vessels with sensory nerves in the joint capsule, anterior disc and retrodiscal tissues. The *left column* shows pgp9.5-positive fibers (all sensory fibers), the *middle column* shows α-SMA-positive (blood vessel) cells and the *right column* shows relative locations of pgp9.5 and α-SMA-positive cells in the mouse joint capsule, articular disc, anterior disc and retrodiscal tissues. Pictures from the joint capsule, articular disc, anterior disc and retrodiscal tissues as well as antibodies used and corresponding colors are indicated. Yellow arrows show pgp9.5^+^ fibers in vicinity of α-SMA^+^ blood vessels, and white arrows show pgp9.5^+^ fibers distanced from α-SMA^+^ blood vessels.

**Figure 12.**
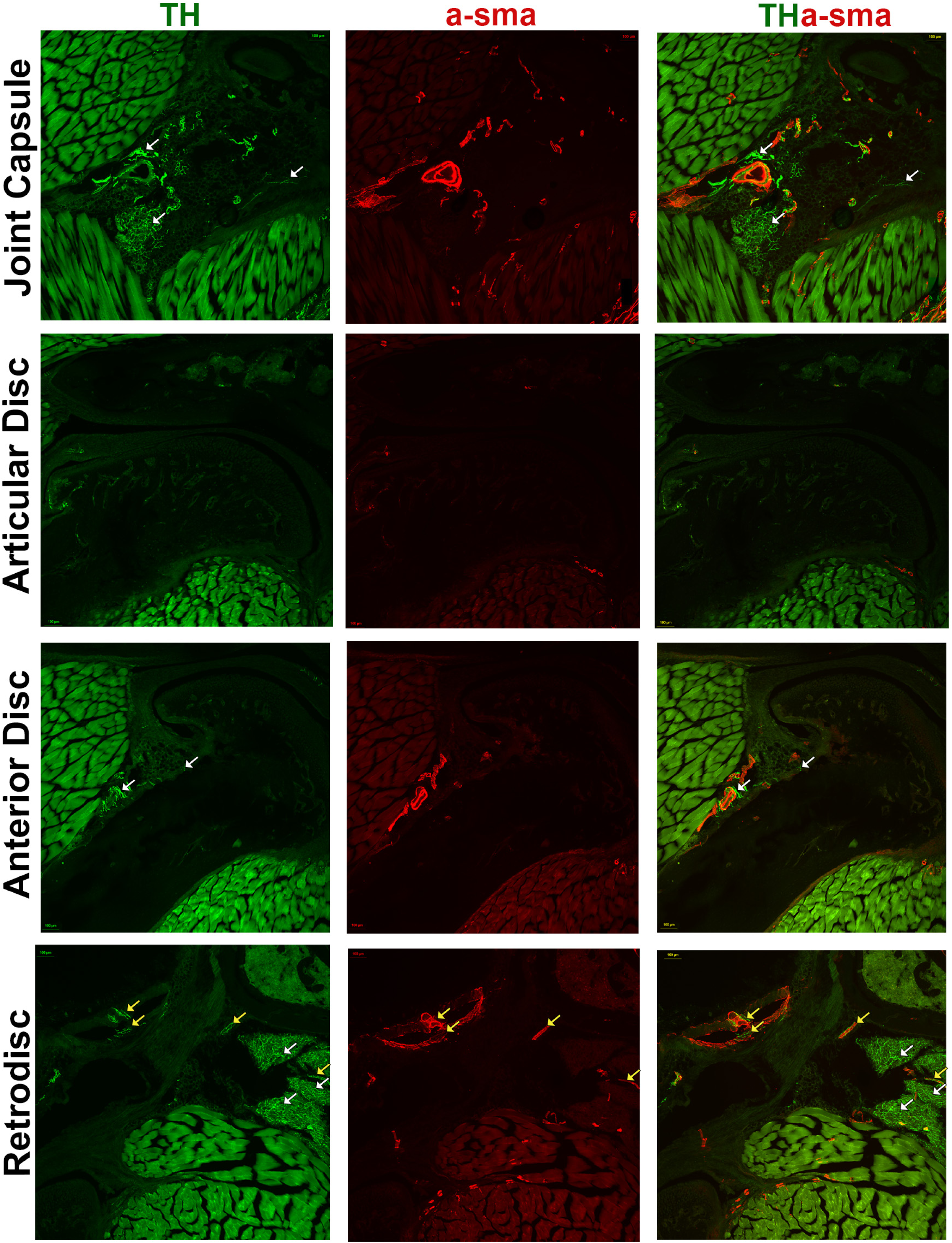
Innervation of blood vessels with sympathetic nerves in the joint capsule, anterior disc and retrodiscal tissues. The *left column* shows TH-positive fibers (sympathetic nerves), the *middle column* shows α-SMA-positive (blood vessel) cells and the *right column* shows relative locations of TH and α-SMA-positive cells in the mouse joint capsule, articular disc, anterior disc and retrodiscal tissues. Pictures from the joint capsule, articular disc, anterior disc and retrodiscal tissues as well as antibodies used and corresponding colors are indicated. Yellow arrows show TH^+^ fibers in vicinity of α-SMA^+^ blood vessels, and white arrows show TH^+^ fibers distanced from α-SMA^+^ blood vessels.

## Discussion

This study aimed to characterize the types of nerves innervating distinct regions of the temporomandibular joint (TMJ), including the joint capsule (with synovium), articular disc, anterior disc, and retrodisc. We found that TMJ tissues are primarily innervated by peptidergic unmyelinated sensory fibers, with the remaining fibers consisting largely of peptidergic myelinated afferents. These results are consistent with previous anatomical and electrophysiological studies identifying various neuropeptides in TMJ-innervating nerves ^7–10^ and classifying them as C– and A-fiber nociceptors ^13,14^. While prior research mainly focused on the joint capsule ^7–10^, our study expanded the anatomical mapping to include the anterior disc and retrodisc. Although the overall innervation pattern was similar across regions, key differences were observed. The retrodisc contained approximately 20% non-peptidergic fibers, including MrgprD⁺, MrgprA3⁺, and MrgprC11⁺ subtypes—fiber types largely absent in other TMJ regions. In contrast, the joint capsule uniquely harbored Sst⁺ fibers. A further distinguishing feature was the expression of Htr3a: nearly all fibers in the joint capsule and anterior disc expressed Htr3a, whereas only about 50% of retrodisc fibers did. These differences suggest that the retrodisc is innervated by a more diverse and molecularly distinct population of sensory neurons.

Electrophysiological studies revealed that TMJ is innervated by C– and A-fiber nociceptors, which could be classified into two main functional subtypes: polymodal nociceptors and high-threshold mechanoreceptors (HTMRs)^13,14^. Building on this, recent integrative analyses of single-cell RNA sequencing data from mouse and human trigeminal ganglia (TG) have identified several molecularly distinct subtypes of peptidergic nociceptors^21^. Among C-fiber nociceptors, these subtypes include Oprk1⁺, Adra2a⁺, Sstr2⁺, and Dcn⁺ neurons. A-fiber nociceptors are categorized primarily as Smr2⁺ and Bmpr1b⁺ neurons ^21^. Both Smr2⁺ and Bmpr1b⁺ A-nociceptors express Htr3a, while only the Oprk1⁺ and Dcn⁺ subtypes among C-fiber nociceptors express this receptor ^21^. In light of our findings, the retrodisc—where only ∼50% of sensory fibers are Htr3a⁺—may contain the full range of peptidergic C– and A-nociceptor subtypes. Conversely, the joint capsule and anterior disc, which show near-complete Htr3a expression among their sensory fibers, are likely innervated mainly by Oprk1⁺ and Dcn⁺ C-fiber nociceptors. These differences point to region-specific specialization in nociceptive innervation within TMJ tissues. A proposed summary of nociceptor subtype distribution across TMJ structures is illustrated in *Figure 13*.

**Figure 13.**
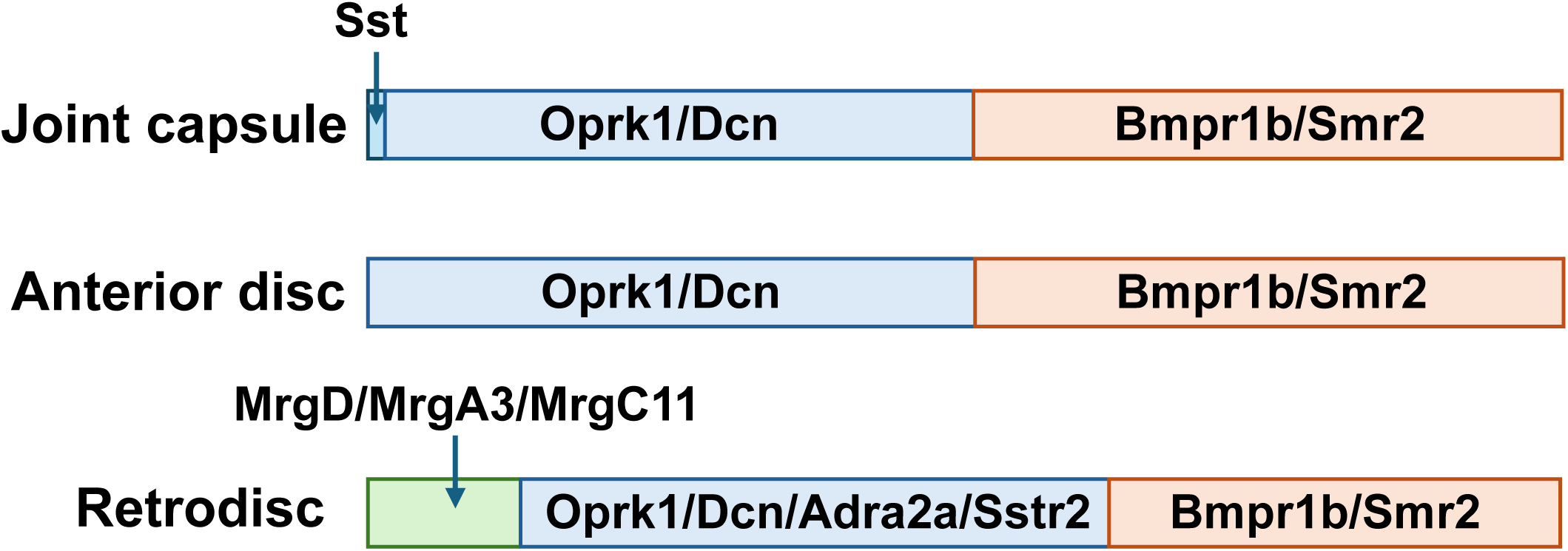
Schematic for sensory neuronal types innervating TMJ tissues. Schematic for sensory neuronal types innervating the joint capsule, anterior disc and retrodiscal tissue. Red box shows myelinated peptidergic nociceptors. Blue box shows unmyelinated peptidergic nociceptors. Green box indicate unmyelinated non-peptidergic nociceptors. Markers for sensory neuronal groups are indicated.

One potential limitation of this study is the exclusive use of male mice. Prior studies utilizing bulk and single-cell RNA sequencing have identified some transcriptomic differences between male and female DRG and TG neurons^12,21,25,26^. Although these differences were generally modest within neuronal populations, notable sex-specific distinctions were observed, particularly in Aβ-rapid adapting (Aβ-RA) and Aδ-low threshold mechanoreceptors (Aδ-LTMRs) in the naïve state^27^. Thus, we do not anticipate sex differences in innervations of TMJ tissues, since they do not contain Aβ-RA and Aδ-LTMRs. Another limitation lies in the manual counting of nerve fibers from IHC images. This method does not distinguish between individual nerve branches and closely adjacent fibers, potentially leading to over– or underestimation. Additionally, we did not use specific markers for neuronal terminals, raising the possibility that some counted fibers were merely passing through the tissue rather than terminating within it. Despite these limitations, the primary aim of the study was to identify predominant and rare nerve fiber types across TMJ tissues. In this regard, the data reliably highlight key patterns of innervation and provide a foundational map of sensory nerve subtypes in the TMJ.

## Materials and Methods

### Ethical approval

This study was conducted in accordance with the ARRIVE 2.0 guidelines ^28^. All animal care and experimental procedures complied with the U.S. Public Health Service Policy on the Use of Laboratory Animals, as well as ethical standards set by the National Institutes of Health (NIH) and the Society for Neuroscience (SfN), with a commitment to minimizing both animal use and suffering. Mice were euthanized via transcardiac perfusion following intramuscular injection of 100 μL of a 1:1 Ketamine/Dexdormitor cocktail. This method, recommended by the AVMA Guidelines for the Euthanasia of Animals, was chosen to ensure minimal distress. All procedures were approved by the Institutional Animal Care and Use Committee (IACUC) of the University of Texas Health Science Center at San Antonio (UTHSCSA) under protocol number 20220069AR.

### Mouse lines

Mice were housed under controlled conditions at approximately 22°C, with 40–60% relative humidity, and maintained on a 12-hour light/dark cycle (lights on at 7:00 AM). Food and water were provided ad libitum in standard home cages. Experiments were performed on adult wild-type C57Bl/6 mice (10–23 weeks old) obtained from The Jackson Laboratory (Bar Harbor, ME).

The following transgenic mouse lines were purchased from The Jackson Laboratory: B6.Cg-Gt(ROSA) 26Sortm9(CAG-tdTomato)Hze/J (Ai9; tdTomato; strain 007909); B6.129P2-Pvalbtm1(cre)Arbr/J (PV^cre^; strain 008069); Mrgprd^tm1.1(cre/ERT2)Wql^/J (MrgprD^cre/ER^; strain 031286), and B6N.Cg-*Sst^tm2.1(cre)Zjh^*/J (Sst^Cre^; stock 013044). The CGRP^cre/ER^ mouse line was generously provided by Dr. Pao-Tien Chuang (University of California, San Francisco), and the MrgprA3^cre^ and MrgprC11^cre/ERT2^ lines by Dr. Liang Han (Georgia Institute of Technology). The Htr3a^cre^ line was developed by the Mutant Mouse Resource & Research Centers (MMRRC) at UC Davis. In inducible Cre-carrying lines, Cre recombination was induced in 6–8-week-old mice via three intraperitoneal injections of tamoxifen (100 mg/kg in corn oil), administered every other day. Recombination was expected to occur within two to three weeks post-injection.

### Tissue collection and processing

All mice were deeply anesthetized via intramuscular injection of 100 μL of a ketamine (75 mg/kg) and dexdomitor (1 mg/kg) cocktail, then transcardially perfused with 25 mL of 4% paraformaldehyde (PFA) in 0.1 M phosphate buffer (PB). Following perfusion, mice were decapitated, and the masseter and temporalis muscles were carefully removed. Bilateral TMJs, along with surrounding bone structures, were then dissected. Facial skin overlying the masseter and TMJ, as well as the TG and dura mater, were collected as positive controls for tdTomato fluorescence in afferent fibers. Dissected TMJs were post-fixed overnight in 4% PFA at 4°C. The next day, tissues were rinsed twice in 1X PB and transferred to 15 mL of 10% ethylenediaminetetraacetic acid (EDTA; ThermoFisher, Cat# 17892) in ddH₂O, adjusted to pH 7.2–7.4. Decalcification in EDTA was carried out over two weeks, with solution changes every three days. After decalcification, tissues were cryoprotected in 10% sucrose for 18 hours, followed by 30% sucrose for another 18 hours. TMJs were then embedded in Neg-50 embedding medium (FisherScientific, Cat# 22-110-617) and cryo-sectioned at a thickness of 20 μm.

### Immunohistochemistry (IHC)

Immunostaining was performed as previously described ^17,29^. Briefly, tissue sections were blocked for 90 minutes at room temperature in a solution containing 4% normal donkey serum (Sigma, St. Louis, MO), 2% bovine gamma-globulin (Sigma-Aldrich, St. Louis, MO), and 0.3% Triton X-100 (Fisher Scientific) in 0.1 M phosphate-buffered saline (PBS). Following blocking, sections were incubated overnight (18 hours) at room temperature with primary antibodies. After incubation, sections were washed with 0.1 M PBS to remove unbound primary antibodies, re-blocked, and then incubated for 90 minutes at room temperature with species-appropriate fluorophore-conjugated secondary antibodies (1:200; Jackson ImmunoResearch, West Grove, PA, USA). Sections were then washed three times for five minutes each with 0.1 M PBS, air-dried, and mounted with Vectashield Antifade Mounting Medium (Vector Laboratories, SKU H-1000-10). The following well-characterized primary antibodies were used on mouse tissue sections: anti-neurofilament heavy chain (NFH) chicken polyclonal antibodies (BioLegend, catalog #PCK-592P, 1:300)^30^; anti-pgp9.5 (Millipore-Sigma, catalog #AB1761-I, 1:400)^31^; anti-CGRP rabbit polyclonal (Sigma, C8198, 1:300)^32–34^; anti-tyrosine hydroxylase (TH) rabbit polyclonal (Pel-Freez; Rogers, AR; P40101; 1:400)^35,36^; and anti-smooth muscle actin (α-SMA) Cy3-conjugated mouse monoclonal antibody (Sigma, C6198, 1:200)^17,37^. Secondary antibodies were generated in donkey (Jackson Immuno-Research).

### Counting of Fibers

Images were acquired using a Keyence BZ-X810 all-in-one microscope (Itasca, IL, USA) or a Nikon AX confocal microscope with Z-stack (“sectioning”) functionality. Most images were captured using a 10× objective. Control IHC was conducted on tissue sections processed identically, but either lacking primary antibodies or both primary and secondary antibodies. Imaging settings were calibrated to ensure that these negative controls produced no detectable nerve fiber signal. Z-stack images for fiber quantification were obtained. For each condition, 3–6 independent tissue sections were analyzed. Sensory fiber quantification was performed manually, following previously established protocols^17^, to estimate the distribution and density of nerve subtypes innervating TMJ tissues. The rationale and validation for manual fiber counting as a reliable method for detecting and quantifying peripheral nerve fibers have been previously described ^17,37^.

### Statistical Analyses

Statistical analyses were performed using GraphPad Prism 10 (GraphPad Software, La Jolla, CA). Data are presented as mean ± standard error of the mean (SEM), with “n” indicating the number of mice analyzed for IHC. For each mouse, fibers were counted from at least 5 randomly selected sections and then the counting was averaged. Group differences were evaluated using appropriate statistical tests, including chi-square analysis with Fisher’s exact test, unpaired t-tests, and one-way or two-way ANOVA followed by Bonferroni’s post hoc test, as applicable. A p-value of < 0.05 was considered statistically significant. Where relevant, interaction F-ratios and corresponding p-values are reported.

## Ethical approval and informed consent

The reporting in the manuscript follows the recommendations in the ARRIVE guidelines (PLoS Bio 8(6), e1000412,2010). All experimental protocols were approved by the UTHSCSA IACUC committee. Protocol numbers is 20220069AR.

## Acknowledgements

We would like to thank Miss. Anahit Hovhannisyan for guidance on IHC. We are grateful to Dr. Pao-Tien Chuang (UC San Francisco, San Francisco, CA) for kindly providing the CGRP^cre-ER^ mouse line; Dr. Liang Han (College of Sciences, Georgia Tech, Atalanta, GA) for kindly providing MrgprA3^cre^ and MrgprC11^cre/ER^ mouse lines generated in Dr. Xinzhong Dong’s laboratory (Johns Hopkins University, Baltimore, MD) and Dr. Liang Han’s laboratory, respectively; and MMRRC at UC Davis for preparing Htr3a^cre^ mouse line.

The RE-JOIN consortium consists of Armen Akopian, Kyle Allen, Alejandro Almarza, Benjamin Arenkiel, Maryam Aslam, Basak Ayaz, Yangjin Bae, Bruna Balbino de Paula, Anita Bandrowski, Mario Danilo Boada, Jacqueline Boccanfuso, Jyl Boline, Dawen Cai, Dellina Lane Carpio, Robert Caudle, Racel Cela, Yong Chen, Rui Chen, Brian Constantinescu, Ibdanelo Cortez, Yenisel Cruz-Almeida, M. Franklin Dolwick, Chris Donnelly, Zelong Dou, Joshua Emrick, Malin Ernberg, Danielle Freburg-Hoffmeister, Jeremy Friedman, Spencer Fullam, Janak Gaire, Akash Gandhi, Terese Geraghty, Benjamin Goolsby, Stacey Greene, Nele Haelterman, Zhiguang Huo, Michael Iadarola, Shingo Ishihara, Azeez Ishola, Sudhish Jayachandran, Zixue Jin, Alisa Johnson, Frank Ko, Zhao Lai, Brendan Lee, Yona Levites, Carolina Leynes, Jun Li, Martin Lotz, Lindsey Macpherson, Tristan Maerz, Camilla Majano, Anne-Marie Malfait, Maryann Martone, Simon Mears, Bella Mehta, Emilie Miley, Rachel Miller, Richard Miller, Michael Newton, Alia Obeidat, Soo Oh, Merissa Olmer, Dana Orange, Miguel Otero, Kevin Otto, Folly Patterson, Marlena Pela, Daniel Perez, Sienna Perry, Theodore Price, Hernan Prieto, Russell Ray, Dongjun Ren, Margarete Ribeiro Dasilva, Alexus Roberts, Elizabeth Ronan, Oscar Ruiz, Shad Smith, Mairobys Soccorro Gonzalez, Kaitlin Southern, Joshua Stover, Michael Strinden, Hannah Swahn, Evelyne Tantry, Sue Tappan, Luis Tovias Sanchez, Cristal Villalba Silva, Airam Vivanco-Estella, Robin Vroman, Joost Wagenaar, Lai Wang, Kim Worley, Joshua Wythe, Jiansen Yan, and Julia Younis.

## Author Contributions

J.J.A.: *methodology, investigation, visualization*. J.J.A. and A.N.A.: *analysis, conceptualization*.

J.J.A. and A.N.A.: *research design* J.J.A., RE-JOIN and A.N.A.– *discussion of results*. J.J.A. and

A.N.A – *drafted the manuscript*; J.J.A., RE-JOIN and A.N.A – *prepared final version of the manuscript*; A.N.A.: *resources, supervision, funding acquisition*. All authors reviewed the manuscript.

## Additional Information

All authors declare that they have no competing interests.

## Legends to Supplementary Figures

**Supplementary Figure 1.**
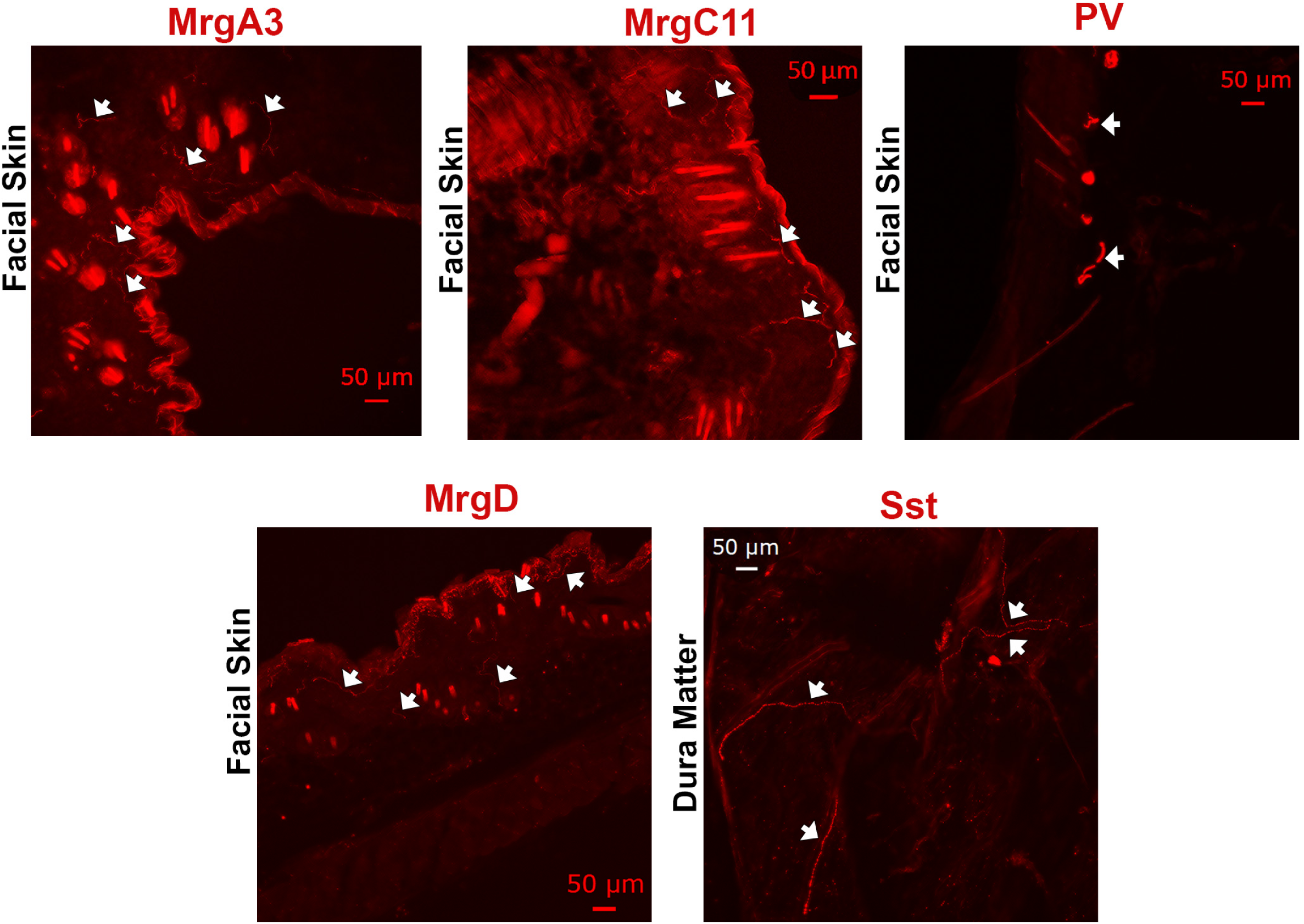
Positive control labeling with MrgprD, MrgprA3, MrgprC11, somatostatin and parvalbumin-positive sensory afferent fibers. *The left panel on the top raw* shows labeling MrgprA3^+^ sensory fibers in facial skin of MrgprA3/tdTom mice. *The center panel on the top raw* shows labeling MrgprC11^+^ sensory fibers in facial skin of MrgprC11/tdTom mice. *The right panel on the top raw* shows labeling parvalbumin (PV)^+^ sensory fibers in facial skin of PV/tdTom mice. *The left panel on the bottom raw* shows labeling MrgprD^+^ sensory fibers in facial skin of MrgprD/tdTom mice. *The right panel on the bottom raw* shows labeling somatostatin (Sst)^+^ sensory fibers in facial skin of Sst/tdTom mice. Markers are indicated panels. White arrows show positive fibers.

